# Subspace reverse-correlation estimation of receptive fields during free viewing

**DOI:** 10.1101/2025.09.24.678404

**Authors:** S. Amin Moosavi, Elaine Tring, Dario L. Ringach

## Abstract

Accurately mapping receptive fields under naturalistic viewing requires correcting for distortions introduced by eye movements. Current approaches often rely on parameterized “shifter networks” optimized with Poisson GLMs, but these methods are computationally intensive and limited to small regions of interest. We present a new framework that exploits the translation properties of Hartley basis functions to model eye position effects directly as phase shifts, eliminating the need for explicit retinal image reconstruction. This formulation admits closed-form gradients, enabling efficient parameter optimization and rapid convergence. Validation on simulated data shows that the method accurately recovers both ground-truth receptive fields and the underlying image transformation. Applied to high-density mouse V1 recordings, the approach improves receptive field sharpness by an average of 56% compared to naive estimates, with optimization completing in minutes on a standard desktop computer. While the method is specific to Hartley basis stimuli, once calibrated, it provides a reusable mapping between eye position and retinal translation. This efficiency and scalability make the technique a practical tool for receptive field mapping in free-viewing experiments and for integration with optical imaging and large-scale electrophysiology.

## 1. Introduction

Accurate estimation of neuronal receptive fields (RFs) in awake, behaving animals is essential for understanding how cortical neurons encode visual information under naturalistic conditions. Eye movements, of course, introduce complications when we are trying to estimate the linear component of a receptive field using reverse-correlation methods [1, 2]. Under these conditions, eye movements continuously shift the retinal image, and the stimulus presented on a screen does not have a fixed relation with the retinal image. If we ignore eye movements, our estimates will be, at best, blurred receptive field estimates and, at worst, they will be just noise. Thus, there have been recent efforts to develop methods that correct for the impact of eye movements on receptive field estimation.

McFarland and colleagues developed an expectation–maximization (EM) framework that couples a linear–nonlinear (LN) encoding model with estimates of eye position to compensate for fixational drifts [3]. By alternating between inferring eye position from V1 population responses and refitting receptive field filters with stimuli corrected for those inferred shifts, they showed that receptive field reconstructions in V1 are markedly improved. Crucially, they demonstrated that eye position itself can be decoded directly from V1 activity with arc-minute precision, surpassing conventional eye-tracking methods [3] (Fig. 1 **a**). More recently, Yates and collaborators introduced a free-viewing, gaze-contingent analysis pipeline that reconstructs the retinal stimulus offline—within a moving ROI aligned to the animal’s eye position, and uses this reconstruction to recover detailed 2D spatiotemporal receptive fields in V1 and MT of freely viewing marmosets (Fig. 1 **b**) [4]. While this paradigm allows one to characterize neural selectivity without fixation training, it relies on high-resolution eye tracking, substantial offline computation for full-field stimulus alignment, and analysis restricted to naturally occurring fixation epochs.

**Figure 1:**
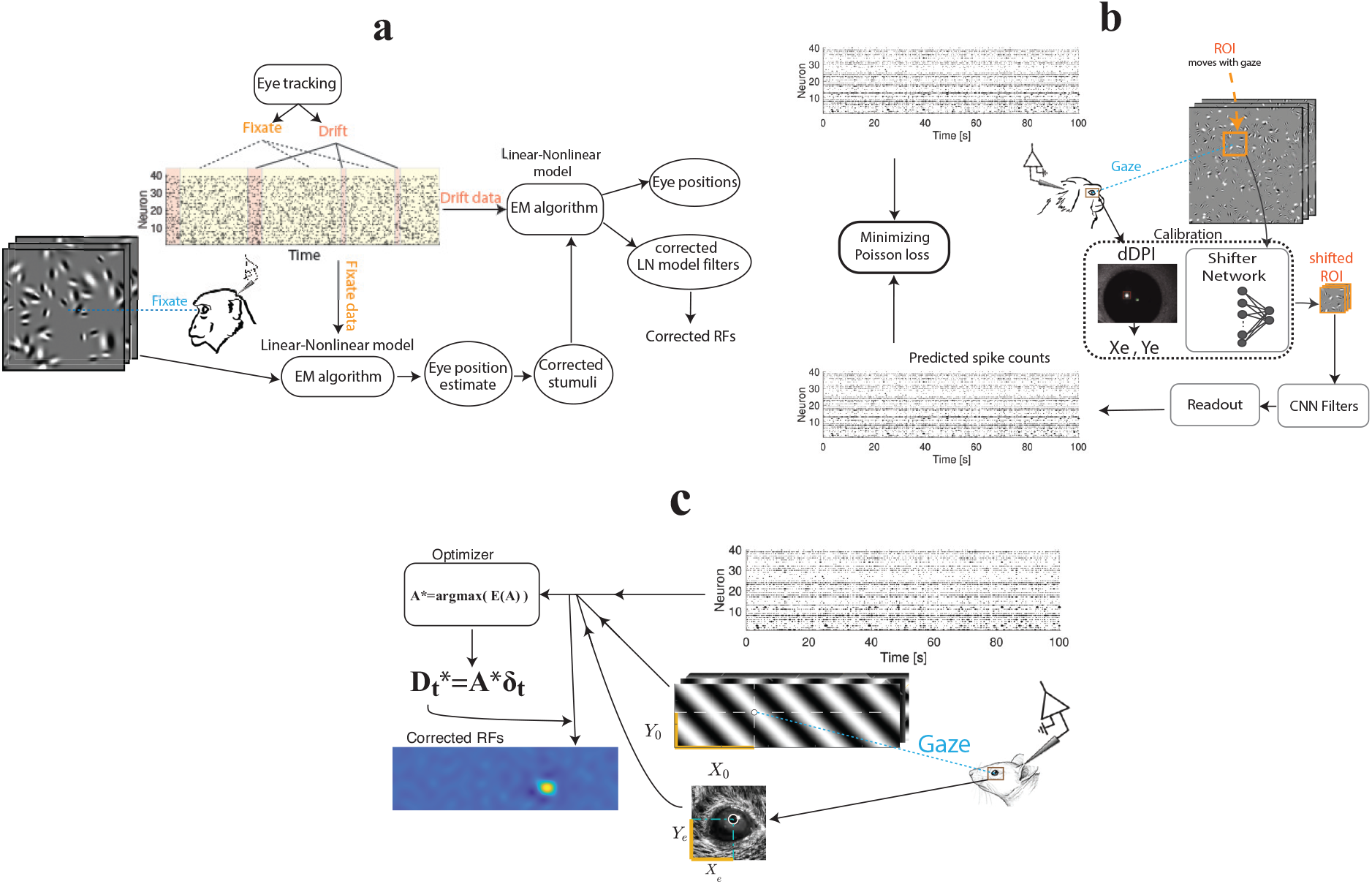
Comparing different approaches to eye movement correction. **a** Diagram of the procedure used by McFarland, et al. [3] in an experiment with a fixation task: While the monkey is instructed to fixate at the screen displaying stimuli, using an eye tracking device, the time is divided into two fixate and drift windows. The eye movement correction is done in two stages. First, the fixating data is used in conjunction with the stimuli to fit a linear-nonlinear (L-N) model of neuronal responses by expectation maximization (EM) algorithm, which estimates the eye positions by which corrections to stimuli are performed. In the second stage, the drift data is used with the corrected stimuli to estimate the eye positions and model parameters, including the L-N filters from which the corrected receptive fields are recovered. **b** Schematics of eye movement correction using a convolutional neural network (CNN) structure utilized by Yates et al. [4]: dDPI devices were used to track the eye position and the corresponding ROI in the visual field that moves with the eyes. Additionally, the system requires calibration, which is performed by shifting the ROIs through a shifter network. The whole network in the system was trained by minimizing the Poisson loss function of the predicted spike counts generated through the CNN. **c** Diagram showing schematics of our method. In our approach, the eye is detected by fitting an ellipse to the pupil at all times. The eye positions are then found by evaluating the center of the pupil in the camera coordinates as (*X*_*e,t*_, *Y*_*e,t*_). We consider the respective gaze position (*X*_0,*t*_, *Y*_0,*t*_). The eye positions are then represented by deviations from their mean value 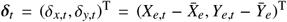 and similarly the gaze position is represented by 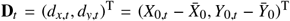. The neuronal spike counts, the stimuli, and the eye position are then fed to the optimizing algorithm maximizing the objective function *E*(**A**). After the optimization, having the optimal 2 × 2 matrix **A**^∗^, using the linear transformation, we evaluate the gaze positions 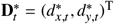 at all times and construct the corrected receptive fields accordingly (See **Methods** for more details).

Here, we introduce an alternative framework for correcting eye movements during receptive field estimation using subspace reverse correlation [5]. The approach requires uncalibrated video recording of eye movements during the experiment. Our optimization procedure operates on the receptive fields of an entire neuronal population distributed across the visual field, rather than being limited to predefined regions of interest. Thus, the method can recover the receptive fields of neurons scattered across the visual field. The algorithm uses gradient descent with a closed-form expression for the gradient, resulting in rapid convergence within minutes on a standard desktop computer.

Briefly, the method works by presenting rapid sequences of sinusoidal gratings (a subset of the two-dimensional Hartley basis set), which uniformly span spatial frequency, orientation, and spatial phase. In our specific application, we record neural responses from the mouse primary visual cortex using high-density silicon probes (Neuropixels 2.0, 4-shank probe), while eye position is simultaneously tracked with a high-resolution camera. Eye position in the camera plane is estimated by fitting an ellipse to the pupil. We model the effect of eye movements as a rigid translation of the image on the retina, with displacements defined as a linear transformation of the eye position. By maximizing the *L*_2_ norm of reconstructed receptive fields, we infer the optimal transformation matrix for eye-movement correction ( Fig. 1 **c**).

After describing the method in detail, we validate it on simulated data, where the ground truth for the receptive fields is known, and show that the technique recovers both the true transformation of the image and provides an accurate reconstruction of the receptive fields. Then, we apply the method to experimental recordings from the mouse primary visual cortex. We find that eye correction leads to a significant enhancement in receptive field signal-to-noise ratios, revealing sharper spatial structures. The method provides a simple and fast alternative for mapping visual receptive fields under free viewing conditions.

## 2. Materials and methods

### 2.1. Experimental model details

All experimental procedures were approved by UCLA’s Office of Animal Research Oversight (the Institutional Animal Care and Use Committee) and were in accord with guidelines on animal research set by the U.S. National Institutes of Health. The data studied here are from a total of 4 mice, male (2) and female (2), aged P35–56, used in experiments where the scientific goal requires an accurate estimation of receptive fields. In other words, the data were not collected for the development of the present method.

### 2.2. Surgery

We performed chronic, extracellular electrophysiological recordings in adult mice (C57BL/6J, both sexes) using Neuropixels 2.0 probes. Each animal underwent a surgery under isoflurane anesthesia (1–2% in oxygen), during which a stainless-steel head bar was implanted, and a small craniotomy ( 1 mm diameter) was made for probe access. A durotomy was performed at the craniotomy site under saline irrigation. Probe insertion was carried out using a Narishige MDS-1 motorized manipulator that controls the insertion speed to minimize dimpling and tissue damage using methods described elsewhere [6]. Recordings were conducted in awake, head-fixed animals positioned on a free-spinning wheel. At the conclusion of the experimental series, animals were euthanized in accordance with AVMA Guidelines. Euthanasia was performed using an overdose of isoflurane followed by cervical dislocation to ensure death.

### 2.3. Visual stimuli

Visual stimuli were presented on a Samsung CHG90 monitor positioned 30 cm in front of the animal. The screen was calibrated using a Spectrascan PR-655 spectro-radiometer (Jadak, Syracuse, NY), with gamma corrections for the red, green, and blue channels applied via a GeForce RTX 2080 Ti graphics card. Hartley functions were generated using a custom-written sketch in Processing 4, leveraging OpenGL shaders (http://processing.org). Hartley functions had the form

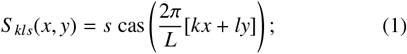

for integer values of −14 ≤ *k, l ≤* 14 and *s* = ±1. Here, the function cas(*θ*) = sin(*θ*)+cos(*θ*), *L* is the size of the screen along the horizontal axis, and (*x, y*) represent a pixel in the screen with those coordinates. Movies were flashed full screen at a rate of 5 fps. Stimulus transitions were signaled to the microscope via a TTL line. As an additional fail-safe, a small square in the corner of the screen flickered at the onset of each stimulus. The flicker was detected by a photodiode, and its signal was also sampled by the microscope.

### 2.4. Data acquisition and pre-processing

Electrophysiological signals were acquired with SpikeGLX [7] and processed using a custom pipeline built on SpikeInterface [8] in Python. Spike sorting was performed with Kilosort [9], and standard quality metrics (including amplitude, isolation distance, contamination index, d-prime, and waveform templates) were logged for all units along with their inferred spatial positions based on probe geometry and template triangulation. SpikeGLX also recorded synchronization TTL pulses from the stimulus computer (stimulus onset/offset), rotary encoder signals from the running wheel, and trigger pulses for two 60 fps behavioral cameras (Allied Vision, U-507m, Stadroda, Germany). One camera captured the image of the contralateral eye, the other a top-down view of the mouse. Infrared illumination and optical filters were used to minimize interference with visual stimulation. Classic image processing, along with Matlab’s function imfindcircles(), was used to estimate the center of the pupil in image coordinates. We then express eye position in each frame as a displacement from the mean position of the eye over the entire experimental session.

### 2.5. Simulation details

In the simulations, we consider *N* neurons with Gabor RFs,

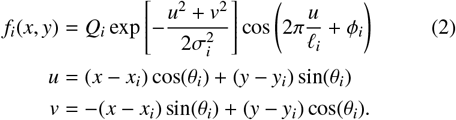

For each neuron *i, Q*_*i*_ is the amplitude, *σ*_*i*_ is the width of the Gaussian envelope, *ℓ*_*i*_ is the period of a sinusoidal carrier, *ϕ*_*i*_ is the spatial phase, *θ*_*i*_ is the preferred orientation, and (*x*_*i*_, *y*_*i*_) specifies the center of the receptive field. We used *Q*_*i*_ = 1, and *σ*_*i*_ = 10 for all neurons, while *ϕ*_*i*_, *θ*_*i*_ were drawn from a uniform distribution in the range [0, *π*], *x*_*i*_ from [0, *L*], *y*_*i*_ from [0, *H*], and *ℓ*_*i*_ from [4*σ*, 8*σ*]. Here, *H* = 135 and *L* = 480 represent the size of the screen (sub-sampled by 8).

The actual eye movements from the experimental datasets are used in the simulations. The firing rate of a neuron *i* in response to a stimulus *S* _*t*_(*x, y*) is a Poisson random variable with rate *λ*_*i,t*_ = [⟨*f*_*i*_(*x* + *d*_*x,t*_, *y* + *d*_*y,t*_)|*S* _*t*_⟩]^+^, where **D**_*t*_ = (*d*_*x,t*_, *d*_*y,t*_)^T^ is the simulated shift in the image caused by the eye movements, brackets indicate the dot product, and [*x*]^+^ represents a half-rectification nonlinearity. This shift of the retinal image is modeled by **D**_*t*_ = **A*δ***_*t*_, where ***δ***_*t*_ = (*δ*_*x,t*_, *δ*_*y,t*_)^T^ is a vector describing the center of the pupil in the two dimensional camera coordinates at trial *t*, with the signals *δ*_*x,t*_ and *δ*_*y,t*_ having zero mean. We adopted a transformation given by

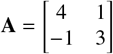

which was selected to be close to the estimated matrix in the experiments (although the results hold for an arbitrary matrix).

## 3. Results

The receptive fields of simple cells in the primary visual cortex of the mammalian brain are well described by two-dimensional Gabor functions, which achieve optimal joint localization in space and frequency domains [10, 11]. The linear component of the response can be efficiently recovered using an orthogonal set of sinusoidal gratings – a subset of Hartley basis functions, *S* _*kls*_(*x, y*) with some maximal spatial frequency (|*k*|,|*l*| *≤ n*_*max*_, see **Methods** for more detail). The result is an estimate of the projection of the receptive field onto the selected subspace [5].

Because of the orthogonality of the stimulus functions (more precisely, they form a “frame”), the receptive field of the *i*^th^ neuron can be estimated as,

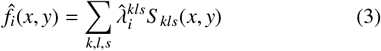

in which 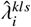 is the estimated firing rate of neuron *i* in response to the stimulus *S* _*kls*_(*x, y*), i.e.

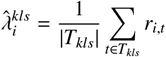

where *T*_*kls*_ is the set of trials with stimulus *S* _*kls*_(*x, y*), |*T*_*kls*_| is the number of trials in the set, and *r*_*i,t*_ is the response of neuron *i* in a trial *t*. In our experiments, the stimuli are balanced, |*T*_*kls*_| = 15.

This naive estimation of the RFs, however, ignores eye movements. When the animal’s gaze shifts, the retinal image of the stimulus is, to a first approximation, translated accordingly. These shifts, which are not accounted for in Eq. 3 cause our estimate to be blurred at best or appear as a combination of multiple copies at different locations in the visual field.

### 3.1. Correction for eye movements

To correct for eye movements, we first used a high-resolution camera to monitor eye position throughout each experiment session. For each frame, we computed the center location of the pupil in camera coordinates (Fig 2**a**). We expressed eye position at each trial as a deviation from its mean across the session, ***δ***_*t*_ = (*δ*_*x,t*_, *δ*_*y,t*_)^T^. How do we find a suitable transformation of the retinal stimulus as a function of eye position? Such transformations, depending on the geometry of the eye and the range of movements, can be expected to be nonlinear. Here, we assumed the range is small enough that a linear approximation would be adequate to compensate for most of the distortion, and modeled the displacement of the retinal image as **D**_*t*_ = **A*δ***_*t*_. Where **A** is a 2 × 2 matrix transforming eye positions ***δ***_*t*_ to Rf displacements **D**_*t*_ = (*d*_*x,t*_, *d*_*y,t*_)^T^.

**Figure 2:**
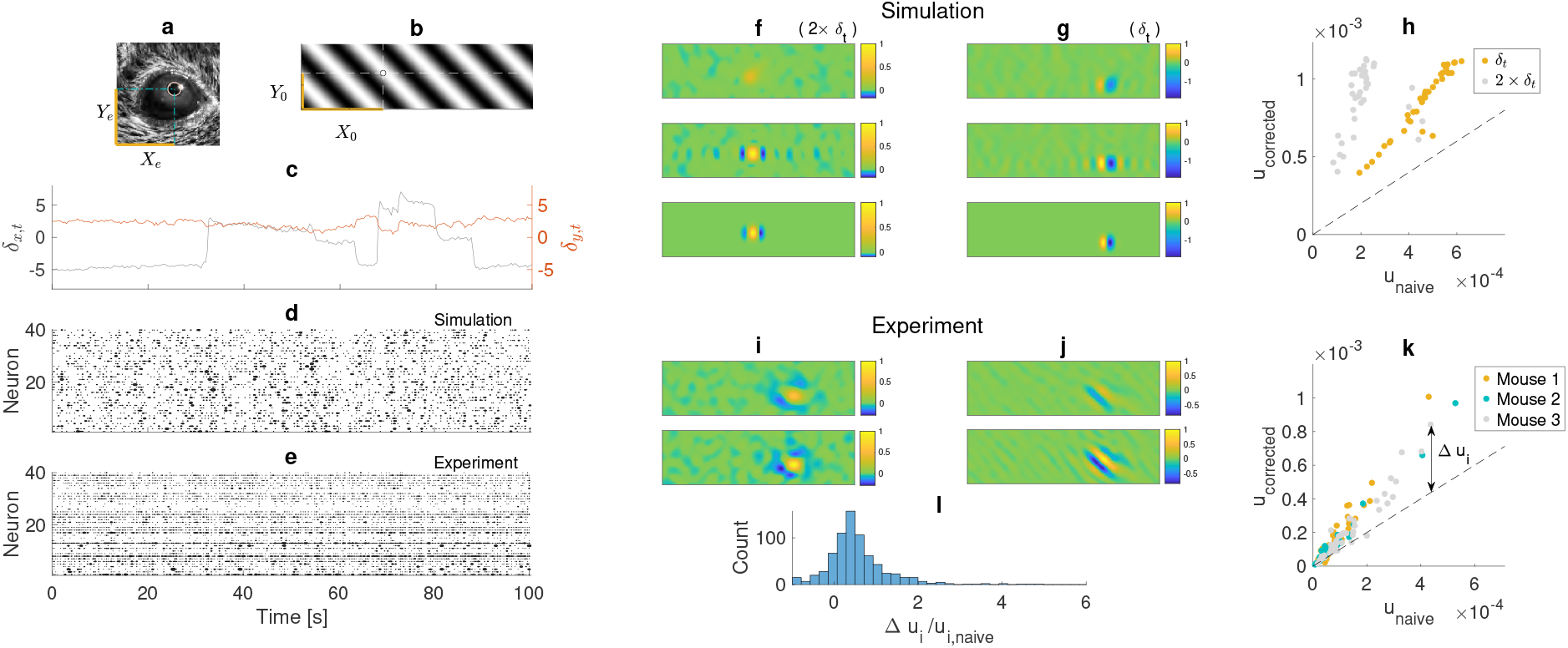
**a** Eye movements were tracked by fitting an ellipse to the pupil (red ellipse). The center of the ellipse is the eye position in the head-centered two-dimensional space of (*X*_*e*_, *Y*_*e*_). **b** A stimulus is shown in the two-dimensional receptive-field space. visual-field-centered coordinates *X*_0_ and *Y*_0_ characterize the gaze location. **c** An example of the eye position time series centered at zero (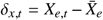 and 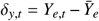). **d** Simulated neuronal responses (40 neurons) are shown over a time window of 100 seconds. Each circle represents the neuronal responses over one trial. The duration of each trial is 200 ms. The size of the circle is a measure of the number of spikes. **e** The same as **d**, but for the experimental data. **f**,**g** Two examples of a simulated neuronal RF are shown. In **f** the simulated eye movements are twice (2 × ***δ***_*t*_) and in **g** equal to that of the experiment (***δ***_*t*_). In both simulations, the top is the naive estimate blurred by eye movements, the middle is the corrected RF, and the bottom is the actual (ground truth) RF used in simulations. **h** Comparing the sharpness of the naive and corrected RFs for simulated data. Colors represent two simulations with domains of eye movements set equal to ***δ***_*t*_ shown in orange, and twice that 2 × ***δ***_*t*_ shown in gray. Each point is for a neuron, and the dashed line is diagonal. Points above the dashed line imply improvement of the RF by eye correction. **i**,**j** Two examples of RF of the mouse V1 neurons. The top is the naive estimate, and the bottom is the eye-corrected RF. **k** RF sharpness improvement for experimental data. Three experiment sessions, one for each mouse, are shown in different colors. **l** Histogram showing the distribution of eye correction improvement over the naive estimate of receptive fields. The data is pooled across 18 experiment sessions with three different mice. For each experiment session, 40 neurons with the most pronounced naive estimates of RFs are selected.

With this model at hand, all we need is to estimate the matrix **A** from the data. As an objective function, we considered the sum of the squared *L*_2_-norm of the estimated receptive fields using

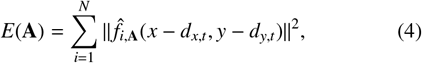

representing the sum of *L*_2_ norms of the receptive fields. In the above equation 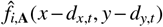 is the translated RF estimate

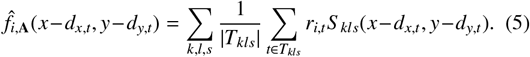

It is not difficult to see why maximizing *E*(**A**) with respect to **A** is an appropriate goal to correct for eye movements. We assume eye position ***δ***_*t*_ is approximately constant during one trial. This is a reasonable assumption because the duration of each trial in the experiments (200 ms) is short compared to the time scale of slow drift eye movements, and saccades will occur on a very small number of trials. Now, if the actual RF of a neuron *i* is *f*_*i*_(**z**) where **z** = (*x, y*)^T^ is a vector in the two-dimensional visual field, for every eye position ***δ***_*t*_, the estimated RF gets displaced to *f*_*i*_(**z** + **D**_*t*_) where **D**_*t*_ = (*d*_*x,t*_, *d*_*y,t*_)^T^ accounts for the RF displacements. Considering *p*(**D**_*t*_) as the probability density function of these displacements across trials, the naive calculation of the RF is then equivalent to the average of all displaced RFs, which in the limit of a long experiment, i.e., a large number of trials, would be:

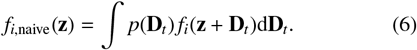

We aimed to correct for eye movements by translating back the RFs according to the linear transformation of the eye position:

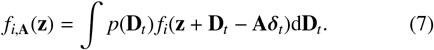

Ideally, **A*δ***_*t*_ = **D**_*t*_ that makes the estimated RF ( *f*_*i*,**A**_(**z**) = *f*_*i*_(**z**)) independent of the eye displacements.

For a population of *N* neurons, the objective function (Eq. 4) can be written as

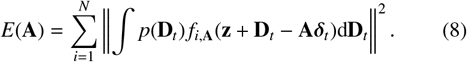

In an ideal eye fixed scenario (*p*(**D**_*t*_) = ***δ***_Dirac_(**D**_*t*_) and ***δ***_*t*_ = **0**) or ideal eye movement correction (**D**_*t*_ = **A*δ***_*t*_) the objective function attains its maximum 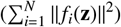, as according to Young’s inequality for convolution [12, 13, 14], therefore

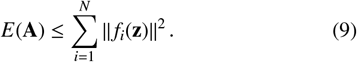

Hence, we define the optimal transformation as the solution to

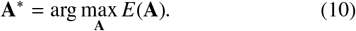

Having **A**^∗^, we corrected the RFs for eye movements by calculating the gaze displacements

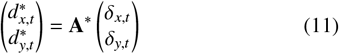

first. Then we obtained the corrected RFs as

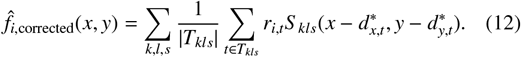

### 3.2. Optimization algorithm

Our optimization algorithm is based on a gradient method. By leveraging the properties of the Hartley functions, we derived an analytical expression for the gradient of the objective function. To do that, we considered the **A** matrix written as

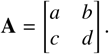

Using **D**_*t*_ = **A*δ***_*t*_ we got

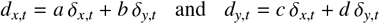

In a four dimensional parameter space (*a, b, c, d*) the gradient of *E*(**A**) = *E*(*a, b, c, d*) is the vector

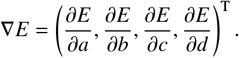

Using Eqs. 1,5 and defining

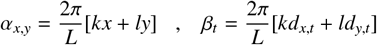

we rewrote the objective function (Eq. 4) as

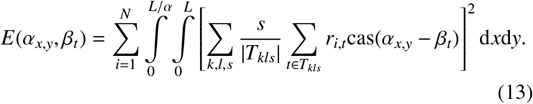

To derive the gradient, we calculated the derivative with respect to the parameters *p* ∈ {*a, b, c, d*} as

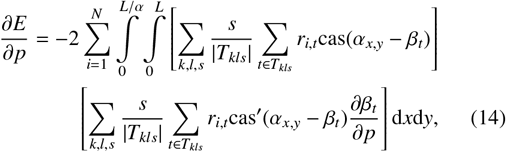

where cas^′^(·) is the derivative of cas(·) and

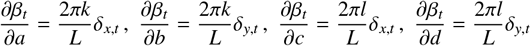

Because the functions cas(*α*_*x,y*_ − *β*_*t*_) and cas^′^(*α*_*x,y*_ − *β*_*t*_) have to be computed at all trials for all the *x, y* domain, their evaluation could, in principle, be computationally expensive. However, the gradient and the objective function computation could be accelerated by decoupling the dependence on the trials *t* and stimulus coordinates (*x, y*). We did so by rewriting the cas(*α*_*x,y*_− *β*_*t*_) and cas^′^(*α*_*x,y*_ − *β*_*t*_) functions as

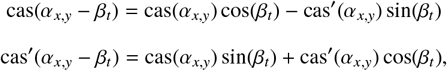

hence, the trial-dependent and (*x, y*)-dependent terms in the square brackets in the above equation got decoupled, e.g.,

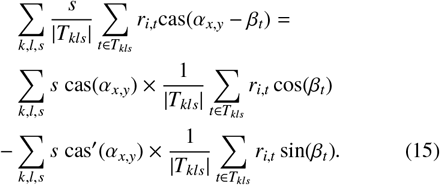

This was particularly helpful because the terms

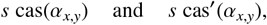

which, due to the dependence on the *x, y* coordinates, were computationally expensive, became independent of the parameters (*a, b, c, d*) and were calculated before starting the optimization algorithm.

Having the objective function and its gradient, we used MATLAB’s fminunc() to find the minimum of −*E*(**A**). Due to the existence of the analytical gradient, the optimization algorithm is efficient, taking only a few minutes to run on a desktop computer.

### 3.3. Validation on simulated data and application to mouse V1 neurons

We first validated the method on simulated datasets where the ground truths, including the matrix **A** and the actual RFs, were known. Gabor filters were selected as the receptive fields of simulated neurons (see **Methods**). Neuronal responses were generated as random samples from Poisson distributions with the rate calculated by an inner product of the RFs and the presented stimuli. An example of the eye position (*δ*_*x,t*_, *δ*_*y,t*_), along with simulated neuronal responses, is shown in Fig 2**c,d**.

Using the simulated dataset (eye positions and neuronal responses), we computed the naive and corrected RFs. Examples of naive RF estimates (Eq. 3) from two simulations are shown in the top panels of Fig 2**f,g**. In mice, eye movements are a fraction of the size of the receptive fields, as opposed to what one expects from recordings in the primate para-fovea. Thus, in our simulations, we considered one case where the amplitude of the eye movements was twice their normal value (Fig 2**f**), and another case where it reflected its normal magnitude (Fig 2**g**). One can appreciate that when the eye movements are large compared to the RF, the naive estimation is substantially blurred and the kernel amplitude is small (Fig 2**f**, top), while the naive estimate is somewhat better when the eye movements are smaller (Fig 2**g**, top). In both cases, the estimated RFs are greatly improved using our correction (Fig 2**f,g**, middle panels) and reflect nicely the structure of the original RFs (Fig 2**f,g**, bottom panels). The reasons the estimates are not perfectly equal to the simulated RFs are due to the fact that we use a finite set of trials and that the recovered RFs represent the projection of the Gabor RFs on the subspace spanned by the set of Hartley functions [5]. In contrast, the matrix **A** was estimated with great accuracy (less than 0.1 percent error).

To further assess the improvements of the RFs, we considered a sharpness measure *u*_*i*_, which for a neuron *i* is defined as the maximum amplitude of the estimated RF envelope divided by its support domain (the domain covering the area around the maximum point with envelope values above 50% of the maximum). The envelope was extracted from the Riesz Transform of the RFs. The sharpness measure of the corrected RFs is plotted against the naive RFs in Fig 2**h**, which shows a significant improvement for all neurons indicated by points above the diagonal dashed line. The improvements were more significant for the simulation with twice-larger eye movements, suggesting that the method would work even better for animals with greater domains of eye movements than mice.

Having established that the algorithm improves the estimation of the simulated RFs, we applied it to the neurons in the mouse primary visual cortex. In our experimental setup, a Neuropixels 4-shank probe was used to simultaneously record the neuronal activities in the mouse V1, giving us the spike times of a few hundred neurons (Fig 2**e**). Using this data, we extracted neuronal responses as the average number of spikes per trial. A high-resolution camera was used to record the eye movements under IR illumination from which we extracted the eye positions (see **Methods**). The eye correction method was applied to the experimental data, similarly to the simulated data. Two examples of naive (top panel) and corrected (bottom panel) RF estimates are shown in Fig 2**i,j**. The sharpness measure of the corrected RF is plotted against that of the naive estimate for three experiment sessions with three different mice in Fig 2**k**. Similar to the simulated data, for most neurons, the points fell above the dashed diagonal line, showing improvements in RF estimates. A summary of the results is shown in Fig 2**l**. The histogram shows the distribution of (Δ*u*_*i*_*/u*_*i*,naive_) with Δ*u*_*i*_ = *u*_*i*,corrected_ − *u*_*i*,naive_, indicating strong improvements across the neuronal populations in 18 experiment sessions with three mice. Overall, the corrected RFs were on average 56% sharper than the naive estimates, and more than 84 percent of neurons’ RFs in all experiment sessions showed improvements.

## 4. Conclusions

In this study, we presented a simple method to estimate the linear component of RFs based on reverse correlation with Hartley basis functions while correcting for eye movements. An efficient algorithm was devised based on the maximization of the squared *L*_2_ norm of the receptive fields of a population using a linear model for the transformation of the retinal image and the fact that rigid translations of the stimulus can be simply represented as changes in its spatial phase. Validation on simulated data confirmed that the algorithm accurately recovers both the underlying transformation and the ground truth RFs in the presence of realistic eye movements. Application of the same method to experimental recordings from mouse V1 showed consistent improvements in RF structure across the population.

Our approach offers a computationally tractable solution that is compatible with large-scale neural recordings and does not require complex probabilistic modeling or machine learning techniques. As a result, the algorithm is computationally efficient, can run on a desktop computer in a few minutes (e.g., the optimization with *N* = 40 neurons converges in less than 3 minutes on a Macintosh machine with Apple M2 Max processor). This efficiency has a price, of course. The method is specific to Hartley basis functions and cannot be easily generalized to other stimulus classes. Still, once the matrix **A** is computed, the system is calibrated in the sense that it provides an estimate of retinal translation as a function of eye position under all experimental conditions. Finally, the model retinal image deformation model is linear, which is adequate for the mouse. More complex models can be implemented within the present framework, provided that gradients with respect to the parameters can be computed.

Although our method was developed in the context of receptive field estimation, the resulting optimal transformation is broadly applicable to other visual areas. This generalization arises from the fact that eye movement–induced displacements are governed by geometric and physical factors independent of brain function. Therefore, once the transformation matrix is estimated, it can be reused to correct for eye movement effects in subsequent experiments involving the same subject and setup. In addition, although the method was tested in simulations and mouse V1 recordings, we do not see any limitations in applying it to other species – in fact, improvements could be even greater in higher mammals, as the eye movements are larger compared to the size of the RFs.

## CRediT authorship contribution statement

D.L.R. and S.A.M conceived and designed the research; E.T. and D.L.R. performed experiments; D.L.R. and S.A.M. analyzed data; D.L.R. and S.A.M. drafted the manuscript; All authors approved the final version of the manuscript.

## Data and Code Availability

A Figshare repository, including data and code, will become available upon acceptance of the manuscript.

